# Cell-autonomous self-renewal

**DOI:** 10.1101/2024.12.20.629654

**Authors:** Ben D MacArthur, Philip Greulich

## Abstract

To self-renew, stem cells must precisely balance proliferation and differentiation. Typically, this is achieved under feedback from the niche; yet many stem cells also possess an intrinsic self-renewal program that allows them to do so autonomously, if required. However, because self-renewal implies a stable equilibrium – in which the expected stem cell number neither increases nor decreases over time – this seems to require fine-tuning to a critical point. Here, we show that this is not the case: self-renewal can, in principle, be easily achieved without the need for extrinsic instruction, feedback or fine-tuning, by a simple cell-autonomous capture-release dimerization cycle that makes use of partitioning errors at cell division to reliably establish asymmetric divisions and perfectly balance symmetric divisions. Such a cycle is seen in the interactions between early-2 factor (E2F) transcription factors cell cyle regulators and their pocket protein partners. Since E2Fs and PPs are also important cell-fate regulators we hypothesise that the E2F-PP cell cycle machinery formed a primordial template upon which self-renewal was first established and continues to be maintained.

## Introduction

Stem cells are present throughout life in most tissues, and are characterized by their ability to self-renew and produce differentiated cellular progeny [1]. Accordingly, they have a central role in regulating tissue growth, turnover, and repair [2]. During development they ensure that cells of all types are appropriately produced and arranged within the context of the growing organism [3]. In healthy adult tissues they regulate homeostatic turnover such that both aberrant growth and tissue loss are avoided, and orchestrate repair as required [4].

Numerous studies have shown that the local microenvironment, or niche, plays a vital part in regulating these abilities [5, 6], by providing chemical [7], mechanical [8] and geometric [9–11] information to the cell. This information is, in turn, processed by intracellular regulatory networks that involve transcriptional regulatory, protein-protein, epigenetic and signalling interactions [12, 13]. Thus, intrinsic and extrinsic regulatory mechanisms collectively ensure that the cell is able to receive environmental cues and act accordingly [14, 15]: for instance, using feedback mechanisms that allow it to sense local cell density or cellular composition and adjust cell division and differentiation propensities accordingly [16–22]. However, to establish self-renewal robustly – that is, in a way that is resilient to inevitable niche fluctuations and insensitive to misguidance due to niche damage, should it occur – stem cells must also possess an internal identity program that safeguards this ability [23]. It has been shown that when detached from their niche, a range of stem cells can maintain self-renewal ability without explicit instruction, indicating the presence of just such a program [24–30]. Yet, the general logic by which it works is not fully known.

In the healthy adult context, stem cell self-renewal can be straightforwardly ensured if each stem cell divides asymmetrically to produce one daughter cell that retains stem cell status and one daughter cell that assumes a differentiated identity, thereby maintaining the stem cell pool at a fixed level, while producing a continuous supply of differentiated cellular progeny [31]. However, from the very earliest stem cell studies in the 1960s onward, it has also been widely observed that stem cell proliferation is an apparently stochastic process that involves balancing symmetric and asymmetric divisions in a probabilistic way [4, 32–36]. In this view, the outcomes of individual cell divisions are driven by stochastic cell-intrinsic processes, which, although they may be biased by signalling from the niche [37], are not themselves closely regulated. Thus, while the asymmetric division strategy implies that individual cell fates are precisely and unambiguously regulated [38], in the stochastic view, self-renewal is established at the level of probabilities, such that cell numbers are appropriately maintained on average within the context of the tissue as a whole [39]. Remarkably, comparison of clone size distributions derived from lineage tracing experiments with those expected from theory [40] suggests that a stochastic model pertains in many adult tissues [41–45], although universality considerations limit the inferences that can be made from clonal data alone [46]. Yet, despite its apparent ubiquity, it is not clear how the precise balance that is needed to implement a stochastic strategy can arise from inherently unreliable stochastic cell intrinsic processes, without some kind of fine-tuning or extrinsic control.

To understand this issue better, consider the following classical model, which views stem cell differentiation as a stochastic branching process [47, 48]:

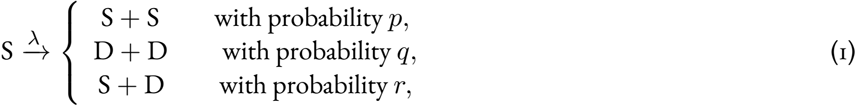

where S denotes a self-renewing stem cell, D denotes a differentiating/differentiated cell, divisions occur stochastically at rate *λ* and *p* + *q* + *r* = 1. Under this model, the expected stem cell number *n*(*t*) evolves as

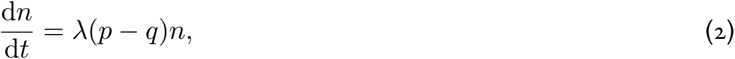

where *f* (*p, q*) = (*p ™ q*) measures cell division bias, which could be niche-regulated.

It is immediate from Eq. (2) that the expected number of stem cells grows if the bias is positive (i.e., if *f* > 0 and S-S divisions predominate over D-D divisions) and declines if the bias is negative (i.e., if *f* < 0 and D-D divisions predominate over S-S divisions). Under a purely asymmetric division strategy this is trivially ensured since *r* = 1 (and therefore *p* = *q* = 0), all divisions are deterministic and the stem cell number is precisely maintained (that is *n*(*t*) = *n*(0) for all *t*). More generally, however, self-renewal can be maintained by the stochastic strategy for *p, q* > 0, so long as *p* = *q*. In this case, cells can divide both symmetrically and asymmetrically and self-renewal is established at the population level, in the sense that while numbers may fluctuate over time, the expected stem cell number remains fixed. However, *p* = *q* is a critical point for the dynamics at which the extinct state (*n* = 0) loses stability. Thus, to maintain a stem cell pool that on average neither grows nor shrinks over time, requires that either the proliferation rate *λ* = 0 or the division propensities are perfectly balanced at this critical point.

Typically, the niche is taken to provide the feedback that is needed to achieve this balance. For instance, most normal cells are subject to contact inhibition [49] and loss of this property is a hallmark of the disregulated growth associated with cancer [50]. The resulting crowding feedback may be easily included in the model above by assuming that the cell proliferation rate *λ* is not constant, but rather depends in a monotonic decreasing way on the total cell number *N*, such that lim_*N*→*K*_ *λ*(*N* ) = 0, where *K* is the tissue’s carrying capacity. Comparable conditions for self-renewal via crowding feedback also hold for more complex cell fate models [16]. In this case, homeostasis is trivially achieved because proliferation ultimately ceases, a situation that is not physiologically realistic. More generally, a state of dynamic homeostasis, in which cell proliferation still occurs but is balanced by stem cell loss (here via D-D divisions), may be achieved by similar regulation of the bias *f* by niche-associated feedback mechanisms [35, 51]. Although the niche clearly provides a collective mechanism by which cell fates can be regulated, it can also be disrupted through injury, chemical damage from toxins, or disease and this, in turn, can lead to macroscopic tissue imbalances, such as hyperplasia and degeneration. If this happens cell-autonomous mechanisms may provide robustness against niche disruption. Besides providing such robustness, primordial cell-autonomous self-renewal mechanisms may also have been important in the initiation of early multi-cellular life, since collective niche regulatory mechanisms had, by definition, not yet been established. This perspective is helpful when looking for possible candidate mediators of cell-intrinsic mechanisms of cell fate balance.

The early-2 factor (E2F) transcription factors and their interaction partners, the Retinoblastoma (Rb) family of pocket proteins (PP), are are highly-conserved and are centrally involved both in cell fate determination and cell cycle regulation [52]. Interestingly, E2F and PP interactions follow a characteristic capture-release dimerization cycle. In G0 and early G1 unphosphorylated PPs form highly stable, transcriptionally repressive complexes with E2Fs [53, 54]. Upon mitogenic stimulation via cyclin, D-CDK4/6 drives PP phosphorylation, which leads to E2F-PP complex dissociation and the release of E2F monomers during mitosis [55]. Moreover, in addition to controlling timely cell cycle progression in essentially all cells, the E2F-PP machinery also plays an important, deeply conserved, part in regulating stem and progenitor cell fate commitment via combinatorial transcriptional regulation of cell fate associated genes [55]. These results suggest that E2F-PP dimerization patterns may have a complex and unappreciated role in intrinsic cell fate balance, based upon their capture-release dynamics. To explore this idea further, here we show how protein dimerization – arguably the simplest and most ubiquitous molecular interaction [56, 57] – can be used to perfectly balance proliferation and differentiation without the need for additional extrinsic instruction, feedback or fine-tuning.

## Results

Motivated by the interactions between E2F and PPs, we consider a pair of proteins A and B that heterodimerize to form an AB complex according to the following reactions:

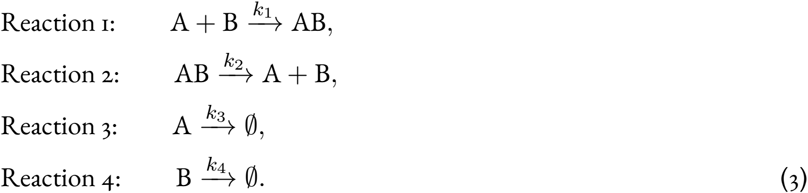

The first two of these reactions describe dimerization and dissociation, while the second two allow for monomer degradation. If all reaction rates are constant and non-zero, then A and B will (in the absence of replenishment) eventually be depleted from the cell and the long-time dynamics are trivial. However, while protein production, interaction and degradation can occur in stable, consistent, ways (e.g., when they regulate vital cellular or physiological processes), their rates are often spatially and/or temporally regulated [58], for instance via reversible post-translational modifications, such as phosphorylation, ubiquitination and glycosylation [59, 60]. The ability of the cell to tune reaction rates is perhaps most important when these reactions participate in the control of dynamic processes such as the cell cycle, which relies on precise, temporally regulated, patterns of protein production and degradation, for instance of cyclins, cyclin-dependent kinases, and other checkpoint regulators [61].

Returning to our earlier example, E2Fs consist of two canonical sub-classes: those that act mainly as transcriptional activators and those that act mainly as repressors [62], although this dichotomy is not rigid [63]. Repressor E2Fs form transcriptional complexes with unphosphorylated PPs in quiescent cells, which suppress the transcription of cell cycle-associated genes and block the G1 to S phase transition. However, on mitogenic signalling, expression of cyclin D-CDK4/6 [64] leads to PP phosphorylation, dissociation of E2F-PP complexes and nuclear export of E2Fs and expression of free activator E2Fs, thereby allowing entrance to S phase. Thus, repressor E2F function is regulated by a ‘dimerization cycle’ in which E2F-PP association occurs in quiescent and early G1 cells, while dimer release occurs during late G1 to M phase. The E2F-PP system therefore provides an important exemplar of how reaction rates are tuned by the cell cycle. However, while much is now known about the molecular details of such dynamics and their variations in different developmental contexts, their general logical consequences have not, to our knowledge, been fully explored. Our purpose here is to show how temporal regulation of dimerization mechanisms could have important consequences for stem cell fate regulation.

Here, we first show how a simple dimerization cycle – that is inspired by E2F-PP dynamics, as a motivating example, yet not intended as a detailed model of this system per se – can make use of stochastic partitioning errors on cell division to reliably establish asymmetric cell divisions, before describing how a generalization of this cycle can be used to precisely balance stochastic symmetric cell divisions, without the need for feedback or fine-tuning of kinetics.

### Generating asymmetric divisions

Typically, asymmetric divisions are regulated by complex mechanisms, that involve both cell intrinsic processes (such as polarization/localization of protein/RNA species, complexes or organelles) and extrinsic processes (such as signaling cues or niche instructions, including mechanical alterations in the local microenvironment) that generate the spatial asymmetry needed to ensure that daughter cells adopt different fates [38, 65]. Here, we outline a simple putative molecular mechanism based on a dimerization cycle that is able, in principle, to reliably produce cells in regulated pairs without the need for active regulatory processes or spatial instruction.

We now consider a scenario akin to that of E2F and PPs, in which the reaction rates in Eq. (3) vary with the cell cycle as follows. (1) During G0 and early G1, dimer dissociation does not occur and the AB dimer – e.g., an E2F-PP dimer – is produced in a stable form, i.e., *k*_2_ = 0; or occurs on a timescale much longer than the cell cycle, i.e., *k*_2_ ≪ 1. In this phase, AB complexes thereby accumulate over time and, since the A and B monomers also degrade a constant rates, all copies of A and B will ultimately be captured in dimer form. These dynamics are illustrated in Fig. 1: we will refer to this as the capture phase of the dimerization cycle and note that it constitutes a simple filtration mechanism that allows the cell to precisely balance expression of A and B prior to cell division. Logically, Reactions 1, 3, and 4 are on during the capture phase and Reaction 2 is off. (2) During late G1-M phase, we assume that AB dimers dissociate – e.g., through the mitogenic stimulation via cyclin D-CDK4/6, which drives PP phosphorylation, forcing complex dissociation and the release of E2F monomers [55]. At this point of the cell cycle, we assume that dimer association and degradation of monomers are inhibited, for instance via the Wnt/STOP stabilization mechanisms that shield most proteins in the late cell cycle from degradation [66,67] (i.e., *k*_1_, *k*_3_, *k*_4_ = 0). In this phase A, B monomers are released into the cell, and, since they do not degrade, they accumulate in precisely equal numbers. These dynamics are also illustrated in Fig. 1: we will refer to this as the release phase of the dimerization cycle. Logically, Reactions 1, 3, and 4 are off during the release phase and Reaction 2 is on. Mathematical details of these dynamics are provided in the **Supplemental Information**.

**Figure 1:**
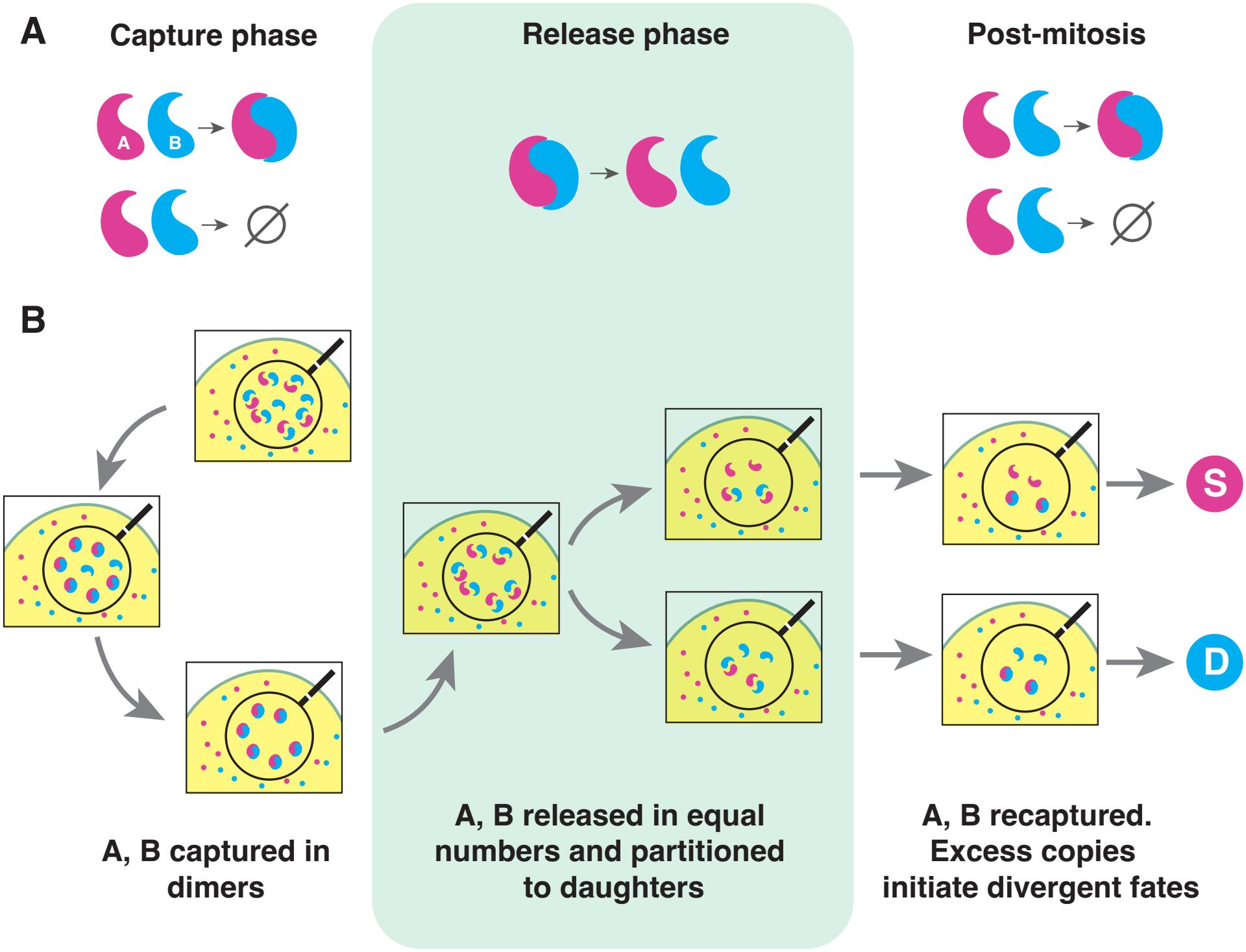
The dimerization cycle. **A, Reactions.** During the capture phase of the cycle (left), two monomers A and B associate to form a stable heterodimer AB and decay at constant rates. During the release phase (middle), dimers dissociate. Post-mitosis, daughter cells re-enter the capture phase (right). **B, Dynamics**. During the capture phase, A and B are sequestered in heterodimers and monomers are cleared from the cell. In the release phase, dimers dissociate and monomers are released in the cell in precisely equal numbers. On cell division monomers are segregated stochastically to daughter cells, which typically results in an excess of A in one daughter and an excess of B in the other. Post-mitosis, A and B are recaptured in dimers and excess copies initiate disparate fates in the two daughters. An asymmetric division is thereby established, which, assuming that A and B determine stem (S) and differentiated (D) cell identities respectively, produces one stem cell and one differentiated cell.

An immediate consequence of this capture-release cycle is that at the point of division A and B are expressed in monomer form in precisely equal numbers in the cell. In principle, the segregation of A and B to daughter cells at this point could depend on many factors, including their spatial localization within the dividing cell, and so could be highly biased and/or correlated. However, regardless of segregation specifics, cell division is an inherently stochastic process and some partitioning errors are expected [68]. Thus, variations in expression of A and B in the daughters will generally occur – i.e., they will not generally both inherit precisely the same number of copies of A and B. Moreover, because the total copy number of each species is conserved, such variations are strictly negatively dependent: if (for example, and without loss of generality) daughter cell 1 obtains A in abundance, then daughter cell 2 must, necessarily, be depleted in expression of A. By extension, if daughter cell 1 inherits more copies of A than B, then daughter cell 2 must, necessarily, inherit more copies of B than A.

This coordinated discrepancy becomes important post-mitosis when, assuming continuity, the daughter cells enter the capture phase of the dimerization cycle. At this point, most free copies of A and B will again be sequestered in AB heterodimers in both daughter cells; yet, after this wave of dimerization is complete, a residual pool of free A will be left in daughter 1 and a residual pool of free B will be left in daughter 2 (or visa versa) in accordance with the stochastic nature of segregation. Thus, the dimerization cycle ensures that daughter cells are always produced in regulated pairs in which one daughter expresses an excess of A in its free form and the other expresses an excess of B in its free form. If A and B are able to drive a cell fate choice – for instance, if they are important transcriptional regulators of the stem cell and differentiated states, respectively – then this discrepancy could provide the impetus needed to establish disparate daughter cell identities in a regulated way. The dimerization cycle is, in principle, therefore a simple way to establish asymmetric divisions, without the need for fine-tuning, spatial feedback, or any other active regulatory process.

Three things should be noted about this cycle. First, the resulting division asymmetry occurs spontaneously, and does not require regulation of segregation dynamics. In the **Supplemental Information** we show that when A and B are segregated purely randomly – i.e., assuming the simplest, most symmetric, model of division in which each individual A or B molecule is assigned to daughter 1 or 2 independently with probability 0.5 – then expression discrepancies between the daughters scale as the square root of the number of molecules of A and B in the parent cell. Thus, even in the least biased model of segregation, if A and B are highly expressed in the parent, then significant differences in expression are expected in the daughters and, since these differences are necessarily negatively correlated, an asymmetric division will ensue.

Second, the resulting division asymmetry does not rely on fine-tuning of any rate constants. The only requirements are that: (1) degradation of A and B monomers during the capture phase occurs sufficiently quickly that all monomers are cleared from the cell before the onset of the release phase; (2) dimer dissociation during the release phase occurs sufficiently quickly that all dimers are dissociated before cytokinesis, and monomers are sufficiently stable that they are not lost during this time; (3) A and B monomers are able to determine the stem cell and differentiated states, respectively, for instance by acting as master transcriptional regulators of these states; (4) the timescale associated with activation of the relevant programs by A and B in the daughter cells is shorter than the timescale associated with monomer degradation. In combination with (1) this means that monomers degrade on a timescale that is longer than that needed to stabilize daughter cell identities, but shorter than the capture phase.

Third, the AB heterodimer does not have a direct regulatory function: it simply acts as a filter that ensures that A and B are expressed in a precise 1:1 ratio prior to cell division. Molecules and reactions that have such indirect functions may be ubiquitous and (as here) vital, yet experimentally underappreciated. We comment on this point further in the **Discussion**.

### Balancing symmetric divisions

In the above, we have considered the dynamics of a single dimerization pair. However, multiple pairs of proteins could undergo similar dynamics; if this happens, then more complex patterns of division can be established. To illustrate, consider the situation in which two sets of proteins – call them A and B, and X and Y – independently undergo the dimerization cycle and stochastic partitioning on division as described above. In this case, if daughter cell 1 inherits more copies of A than B and more copies of X than Y (for example, and without loss of generality), then daughter cell 2 must necessarily inherit more copies of B than A and more copies of Y than X. After mitosis, when the daughter cells enter the capture phase of the dimerization cycle, most copies of A and B will, as before, be sequestered in AB dimers and most copies of X and Y will be sequestered in XY dimers, leaving a residual pool of free A and X in daughter 1 and free B and Y in daughter 2. Thus, daughter cells are again produced in regulated pairs. However, while only one outcome is possible when only one dimerization pair is involved, and the division is necessarily asymmetric, here two outcomes are possible: if daughter 1 expresses free A and X, then daughter 2 must express free B and Y; if daughter 1 expresses free A and Y, then daughter 2 must express free B and X. The contingency table associated with these divisions is shown in Fig. 2A. Moreover, if each pair is segregated independently – i.e., if no additional regulatory mechanisms that coordinate their distribution to daughters are involved (mathematically, if *P* (*a, b, x, y*) = *P* (*a, b*)*P* (*x, y*), where *P* (*a, b, x, y*) is the joint distribution of A, B, X and Y in a daughter cell and *P* (*a, b*) and *P* (*x, y*) are the marginal distributions) – then both of these outcomes are equally likely, regardless of any other segregation specifics: that is, they both occur with probability 0.5. Thus, while a single dimerization pair can robustly generate asymmetric divisions, two dimerization pairs can perfectly balance symmetric divisions. These possibilities are shown schematically in Fig. 2B.

**Figure 2:**
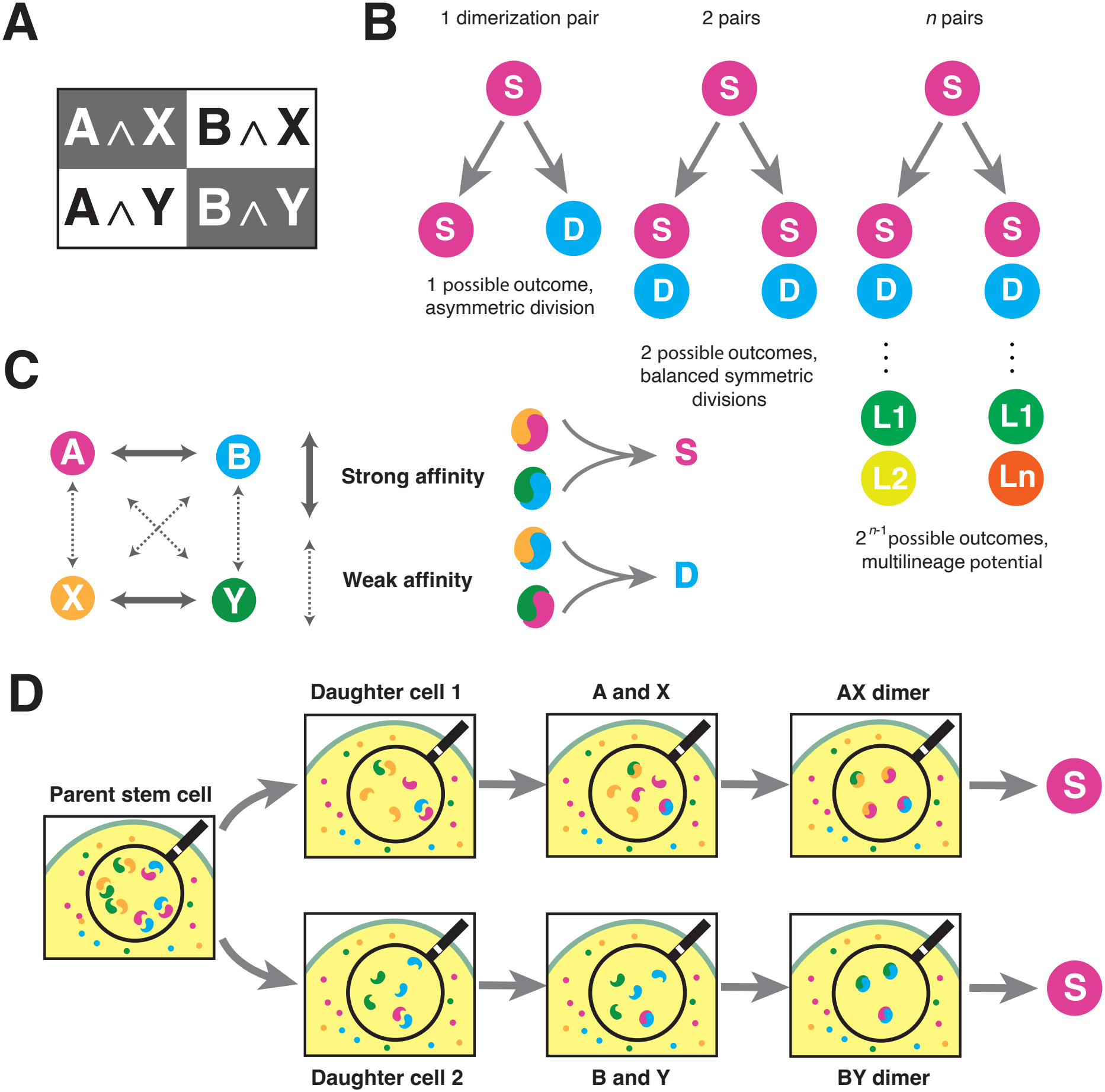
Dimerization dynamics in higher dimensions. **A, 2 *×* 2 contingency table.** When two dimerization pairs are present, daughter cells are produced in regulated pairs (here, shown in gray and white), that each occur with probability 0.5. **B, canonical types of division**. If cell identities are determined by one dimerization pair then all divisions are asymmetric (left). Assuming one protein in the pair determines the stem (S) cell identity and the other a differentiated (D) cell identity then an S-D division ensues. If identities are determined by two dimerization pairs, then two types of division are possible (middle), according to the contingency table in panel **A**. If interactions between the proteins establish an AND logic (see panels **C-D**, and Fig. 3), then S-S and D-D divisions occur in precise balance. If identities are determined by *n* dimerization pairs then 2^*n*−1^ types of division are possible, each occurring with probability 2^1−*n*^. Daughter cell fates are then determined by secondary interactions between the dimerizing proteins, and/or other regulatory mechanisms, which provide the potential to precisely balance the production of cells from multiple lineages (here, labeled L1, L2, … Ln). **C, example of a competitive dimerization network** that is able to balance symmetric divisions; AX and BY dimers determine the stem cell state and BX and AY dimers determine a differentiated state. **D, schematic of illustrative dynamics**. On division, daughter cell 1 inherits A and X in excess. Daughter 2 therefore inherits B and Y in excess. AB and XY dimers form preferentially according to the logic in panel **C**. Residual copies of A and X in daughter 1 then form AX dimers; similarly, residual copies of B and Y in daughter 2 form BY dimers and an S-S division ensues. By similar reasoning, if daughter cell 1 inherits A and Y in abundance, then daughter cell 2 must inherit B and X in abundance (not shown). AY and BX dimers then form and a D-D division ensues, again according to the logic in panel **C**.

In the context of stem cell proliferation a single dimerization pair can support self-renewal via asymmetric divisions if A and B are determinants of the stem cell and differentiated cell states, respectively (or visa versa). Similarly, two dimerization pairs can support self-renewal by balancing symmetric S-S and D-D divisions if (A and X) or (B and Y) determine the stem cell state, and (A and Y) or (B and X) determine the differentiated state (or visa versa).

The required and logic could be achieved numerous ways. For example, A and B may bind to X and Y with weak affinity, either exclusively or with other co-factors, to form AX and BY or AY and BX transcriptional complexes that act as master regulators for the stem cell and differentiated states, respectively. Since they are formed with weak binding affinity, such secondary complexes would only generally be produced from the residual monomers left once the primary AB and XY complexes have saturated post-mitosis. This mechanism is an example of a competitive dimerization network, and is illustrated in Fig. 2C-D. Such networks are known to be highly expressive and easily implemented, and therefore may be widely used by the cell to efficiently regulate a host of cell behaviours [69]. Indeed, the E2F-PP network is characterized by such competition. For example, E2F1, E2F2 and E2F3 bind preferentially to Rb, while E2F4 and E2F5 bind only weakly to Rb; similarly, E2F4 and E2F5 bind preferentially to p107 and p130, which have weaker affinities for E2F1, E2F2 and E2F3 than Rb [52]. This analogy is supported by the known ability of such cross-dimers to affect cell fate choice. For instance, in adult epidermal progenitors, expressing synthetic hybrid constructs with interchanged pocket-protein binding regions (E2F1/4 and E2F4/1) severely impairs keratinocyte differentiation and suppresses colony-forming efficiency, indicating that shifting the exact E2F-PP binding interface fundamentally disrupts self-renewal [70]. Similarly, in adult neural stem cells the non-canonical p107/E2F complex serves to inhibit symmetric expansion; p107-deficient neural precursors demonstrate striking, cell-autonomous up-regulation of Notch-Hes1 signaling and an enhanced capacity for self-renewal [71].

As a alternative mechanism, one pair – A and B, say – may act as pioneer factors [72] that initiate chromatin opening and allow transcriptional regulation of genes associated with stem cell and differentiation programs (call them the S- and D-programs), respectively. If X acts as a ubiquitous transcriptional activator and Y as a repressor, then a similar and logic can arise via temporal regulation of gene expression, assuming that both the S- and D-programs will eventually open post-mitosis (i.e., independently of expression of A and B) and they mutually repress each other when active. Such logic has been widely implicated in directing cell fate and reprogramming by establishing molecular switches that stabilize nascent commitment events [73–76], which in turn activate downstream regulatory networks [77]. In this case, two general possibilities can occur: if either A or B is co-expressed with X in a daughter cell, then the respective program will be opened early and activated, thereby allowing it to establish early dominance. Conversely, if either A or B is co-expressed with Y in a daughter cell, then the respective program will be opened early and repressed. In this case, assuming that DNA-bound Y is stable and free Y is cleared from the cell on an intermediate timescale, once the opposing program opens it will establish dominance over the repressed program via constitutive mechanisms. This mechanism is illustrated in Fig. 3. Such interactions between transcriptional regulatory mechanisms and chromatin remodelling agents are known to be important E2F-PP pathways, for instance to generate context-specific patterns of DNA association [78, 79].

**Figure 3:**
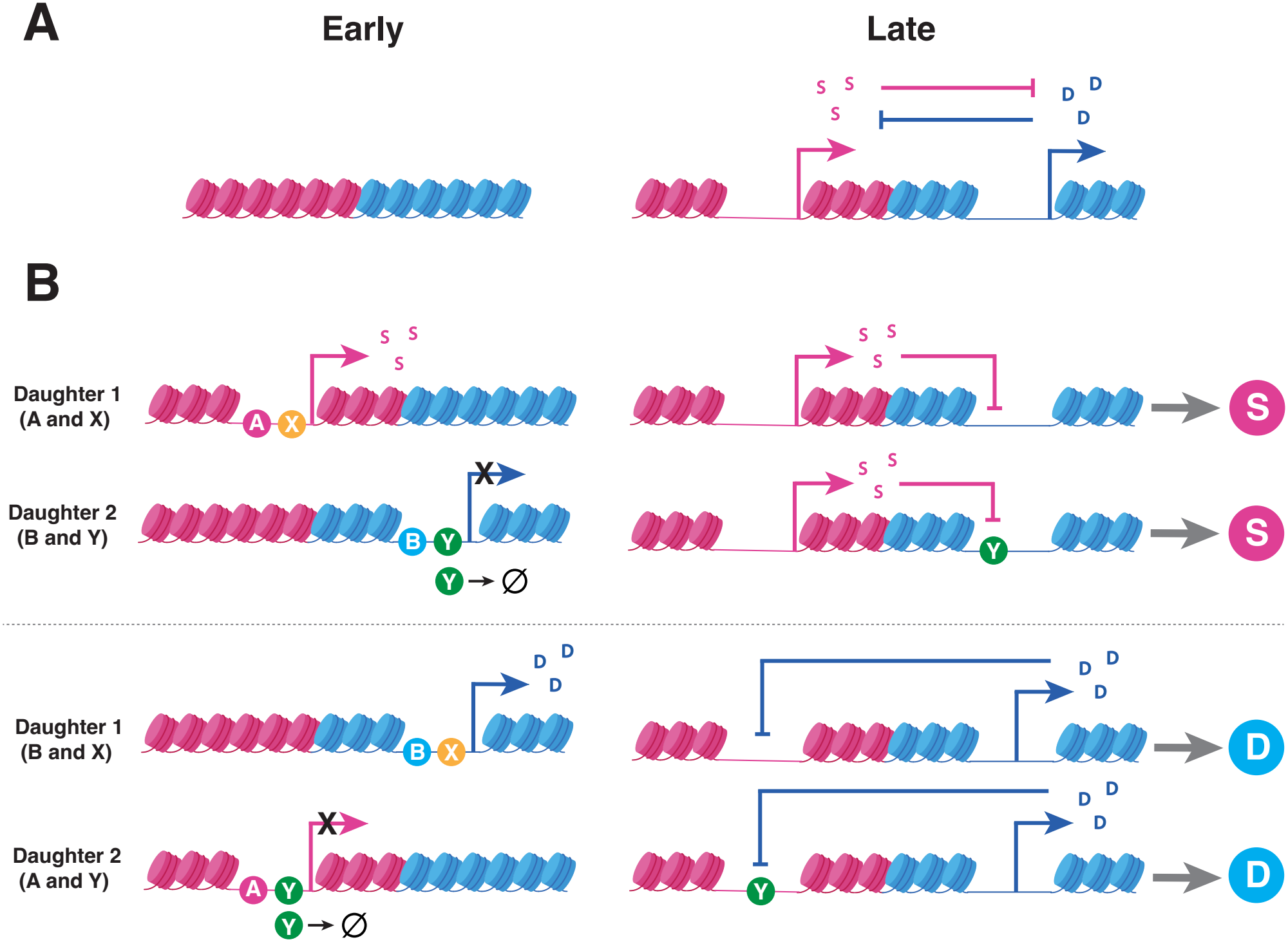
Epigenetic regulation of daughter cell fates. **A, expression logic.** Each daughter cell has the capacity to express a stem cell program (pink) and a differentiation program (blue). In the absence of any regulatory factors, chromatin is packed and both programs are closed early post-mitosis (left), and both open on a slow timescale (right). Once active, these programs interact via mutual repression (right). **B, Dynamics post-mitosis**. In the presence of A and B the respective programs can be opened early. (First row) If A and X are expressed in daughter 1, then A opens the S-program early which X activates (left), allowing it to establish dominance over the D-program once it opens (right). The first daughter cell then adopts a stem cell identity. (Second row) B and Y are expressed in daughter 2, according to the logic in Fig. 2A. B opens the D-program early, which Y represses (left). Assuming that free copies of Y monomers are cleared from the cell by degradation, and DNA-bound copies are stable, the S-program is then opened at a later time and establishes dominance via constitutive mechanisms. The second daughter then adopts a stem cell identity, and an S-S division ensues. (Third and fourth rows) by a similar reasoning, co-expression of B and X in daughter 1, and A and Y in daughter 2 gives rise to a D-D division. Adapted from [80].

## Discussion

To self-renew stem cells must be able to perfectly balance proliferation and differentiation. Here, we have shown how a simple dimerization cycle can, in principle, achieve this balance without feedback or fine-tuning. This mechanism does not replace proven niche regulatory processes, but may provide a basic layer of control that safeguards self-renewal against transient environmental fluctuations, conflicting or unreliable niche instructions.

We have presented the basic logic of the dimerisation cycle in a deliberately abstract way in order to clarify its consequences. The specific nature of the proteins involved is still to be determined. Because tissue homeostasis requires the presence of a self-renewing sub-population of cells [36], the establishment of self-renewal is a pre-requisite for complex multicellular life that conceivably first arose from modifications to cell cycle machinery, before being adapted by niche regulatory mechanisms. We speculate that due to its emergence in early eukaryotic evolution [81] and since it plays a deeply conserved part in regulating both cell cycle progression and stem cell fate commitment throughout the animal kingdom [55], the dimerisation logic encoded in the E2F-PP system may have provided a template on which self-renewal capability was first established and continues to be maintained. This hypothesis is amenable to experimental verification. For instance quantitative live-cell imaging techniques, such as Förster resonance energy transfer (FRET), could be used to track the real-time stoichiometry of proteins, to confirm the 1-to-1 ratios prior to cell division, and transient cross-dimerization of variants immediately following cytokinesis. Additionally, CRISPR/Cas9 mutagenesis or targeted small-molecule blocking of relevant dimerization reactions [82] could be used to produce variants with a lower dimerization rate, in order to synthetically flatten the specific binding hierarchies without completely disrupting protein function. If the proposed mechanism holds, then short-circuiting this post-mitotic cross-dimerization logic would purposefully disrupt the balance of symmetric divisions and manifest as a quantifiable loss of tissue homeostasis in stem cell lineage tracing assays.

Based on this mechanism we propose two canonical modes of stem cell division: (M1) if divisions are exclusively regulated by one dimerization pair, then asymmetric divisions will ensue; (M2) if divisions are regulated by two dimerization pairs, then perfectly balanced symmetric divisions will ensue. (If divisions are regulated by more than two dimerization pairs then these canonical modes can be generalized to allow multi-lineage potential, see Fig. 2B.) However, in general, stem cells are able to divide both asymmetrically and symmetrically [4] and these two modes of division are not mutually exclusive. Indeed, they may be easily integrated by assuming that the stem cell divides asymmetrically by default, and symmetric divisions only occur if the impetus to do so is sufficiently strong. This may occur, for instance, only if the expression discrepancies associated with symmetric divisions exceed some threshold that reflects niche ‘instructiveness’. In circumstances that require precisely instructed asymmetric divisions (i.e., when stem cell number fluctuations cannot be tolerated) this threshold will be high and all, or almost all, divisions will be asymmetric. In other circumstances, the niche may not need to exert such direct control, and the threshold will, accordingly, be low. In such cases, balanced symmetric divisions will predominate. At intermediate threshold levels, both division modes will be active, and symmetric S-S and D-D divisions will occur equiprobably at a niche-defined rate. Mathematical details of this integrated model are provided in the **Supplemental Information**.

To illustrate these canonical modes of division, we have focused on their potential role in regulating the classical stochastic self-renewal strategy given in Eqs. (1)-(2). However, other stochastic strategies are possible. For example, it has been observed that many stem cell populations exist in a state of dynamic heterogeneity [83, 84], in which individual cells transition stochastically between multipotent, primed and differentiated states over time [85,86]. Although unpredictable at the single cell level, this switching establishes a stable, structured population that is able to adapt to a variety of environmental changes in a maximally responsive way [39, 87, 88]. Because it provides a way for robust functional properties to emerge at the macroscopic (i.e., population) level from inherently stochastic microscopic (i.e, cellular) dynamics this is an attractive perspective [89]. Notably, the standard model of stem cell proliferation based on such dynamic heterogeneity assumes that stem cell divisions are all asymmetric by default, and symmetric divisions arise from stochastic cell state switching subsequent to division [35]. Our framework is therefore consistent with this view, although in our model division asymmetry arises as a consequence of the dimerization cycle and stochastic partitioning of cell fate determinants on division, rather than being assumed. More generally, our framework provides a mechanistic basis for the underpinning assumption that daughter cell identities are not determined prior to mitosis, but rather arise from regulated variations in initial conditions, which initiate disparate stochastic cell fate trajectories in the daughters post-mitosis.

In this regard, it is notable that stochasticity has a vital role in our framework. In particular, we do not make any assumptions concerning the segregation dynamics that occur at mitosis except that: (1) some variation in expression in the daughters is required and (2) if multiple dimerization pairs are involved then each pair is segregated independently of the others. Thus, partitioning errors, which can be significant due to upstream stochastic effects, daughter cell size discrepancies and/or spatial clustering of proteins [68], are welcome. In principle, ordered segregation mechanisms, which reduce such errors are possible, although they are harder to implement [68]. In this case, they would also be counterproductive. Thus, our framework requires minimal regulatory control and is robust to segregation specifics and, in fact, benefits from the disorder that is inherent to partitioning of cellular contents on division. As such it provides another example of how cellular systems can make good use of noise [90–92].

Although we have presented the framework in a general way, it is consistent with well-known properties of cell fate regulatory networks. For example, it has been widely observed that cell identities are controlled by small sets of master lineage specifiers that co-regulate diverse genomic targets [93, 94], and identities can be reliably guided by controlling these sets [95–98]. Moreover, these factors often associate with each other, either directly or via co-binding to third factors [99–102], giving rise to complex regulatory networks that involve both protein-DNA and protein-protein interactions. However, despite the apparent complexity of these networks, the dimerization mechanism we propose is remarkably simple and does not depend on complicated regulatory processes or feedback control for its efficacy. As such, it is consistent with a continuous view of cell fate commitment [103–105]. This simplicity is notable for two reasons. First, it suggests that important aspects of self-renewal may be essentially low-dimensional and therefore amenable to intervention. This underscores the importance of simple molecular processes such as dimerization as therapeutic targets [56, 69]. Moreover, because dimerization serves primarily as a means to balance the expression of key cell fate determinants rather than produce a functional biomolecule per se, it suggests that important aspects of stem cell identity may be regulated in indirect ways. We conjecture that aspects of cell behaviour in other contexts are similarly regulated. Second, it provides a particular justification for why dimerization is such a ubiquitous bio-chemical reaction [57]: it is a very simple way to regulate the co-expression of competing factors, and thereby precisely balance their downstream effects, without the need for feedback or fine-tuning of reaction rates. Thus, it provides an effective way for cells to hedge their bets in the face of environmental unpredictability [87]. As such, dimerization-based mechanisms may be evolutionarily favoured whenever cells must act under conditions of uncertainty. Although we have focused on stem cell self-renewal, decision-making under conditions of uncertainty is a challenge faced by cells throughout nature, in both unicellular and multicellular contexts [106]. The mechanism described here, or variations thereof, may therefore be relevant elsewhere.

## Supporting Information

### Reaction rate equations and stability analysis

The dynamics of the system of reactions given in Eq. (3) are described by the following set of reaction rate equations:

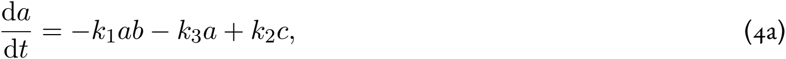

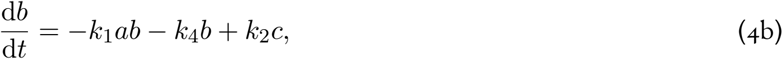

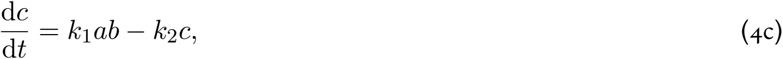

where lower case letters denote concentrations, and *c* denotes the concentration of the AB dimer.

In the capture phase of the dynamics, dimer dissociation is off and *k*_2_ = 0. In this case, the equations for the steady state are

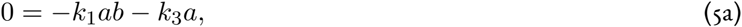

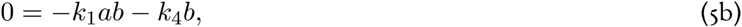

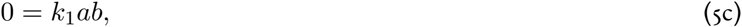

which have only one solution: *a* = *b* = 0 and *c* = *c*_ss_ for some *c*_ss_ > 0 (assuming *a*(0), *b*(0) > 0). The Jacobian matrix of the system Eqs. (4a)-(4c) at this steady state is,

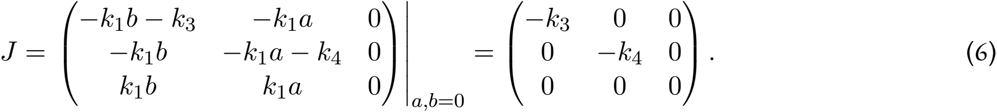

The eigenvalues of this matrix are (0, *™k*_3_, *™k*_4_). The zero-eigenvalue corresponds to perturbations in the manifold *c* = *c*_ss_, which are determined by initial conditions. Stability to perturbations off this manifold (that is, for any fixed value of *c*_ss_) is determined by the remaining eigenvalues, which are both negative. Hence this steady state is stable, and is a universal attractor for the dynamics. Thus, as the end of the capture phase approaches, *a, b* → 0 and *c* → *c*_ss_ > 0. The expected copy number of the AB heterodimer therefore approaches *C*_ss_ = *V c*_ss_ where *V* is the cell volume, and the expected number of A, B monomers both approach zero.

In the release phase, all these dimers dissociate. In this phase, *a* − *b* is a conservation law for the dynamics and so *a*(*t*) = *b*(*t*) for all *t*. The expected number of both A and B monomers will therefore tend to *C*_ss_ as the end of the release phase approaches. Thus, they will be present in equal numbers, regardless of rate parameters in Eqs. (4a)-(4c).

### Partitioning errors

Consider the expression levels of a pair of proteins A and B, with dynamics as described in the main text, in a stem cell under-going mitosis. Let *m* denote the copy number of A (and therefore B) in the cell at this time. Assume that on division both A and B are partitioned independently and equiprobably in the daughter cells, i.e., each individual A or B molecule is assigned to daughter 1 or 2 independently with probability 0.5. Both A and B will then be Binomially distributed in the daughter cells immediately after division. Specifically,

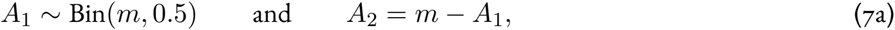

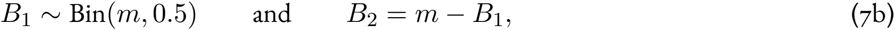

where *A*_*i*_ and *B*_*i*_ denote copy numbers of A and B in daughter *i* respectively. Assuming that the parent stem cell contains an abundance of A and B (i.e., *m* is large) we may take the Normal approximation to the Binomial. In this continuum limit,

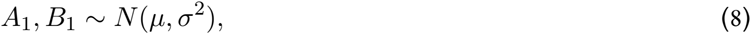

where *µ* = *m/*2 and *σ*^2^ = *m/*4.

To understand how variations in the expression of A and B generate asymmetric divisions, we are concerned with differences in their expression and particularly with the distribution of |*A* − *B*| in the daughter cells (note that by symmetry |*A*_1_ − *B*_1_| = |*A*_2_ − *B*_2_|). For *m* large, it is immediate that

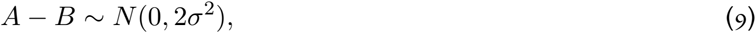

in both daughters and so |*A* − *B*| follows a folded Normal distribution with mean zero, and variance 2*σ*^2^ (here, we have overloaded notation to let |*A* − *B*| denote the absolute difference in copy number of A and B). Thus,

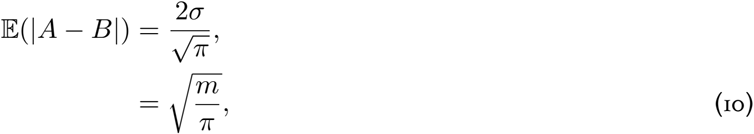

and the expected difference in expression of A and B in the daughter cells scales as the square root of their copy number in the parent stem cell.

### Integrating asymmetric and symmetric divisions

If, in addition to A and B, a second pair of proteins, X and Y, independently undergo a similar dimerization cycle, then

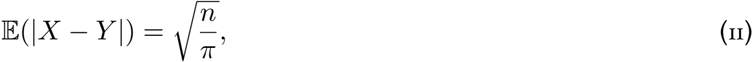

where *n* is the copy number of X (and therefore Y) in the parent cell prior to mitosis. Assuming that the stem cell undergoes a symmetric division only when |*A ™ B*| and |*X ™ Y*| both exceed a threshold *T* in the daughters – which, for instance, ensures that the secondary AX, AY, BX and BY dimers described in Fig. 2C are produced in sufficient numbers – the probability that a symmetric division occurs is given by

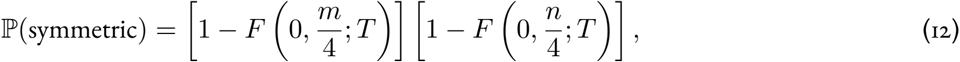

where *F* is the cumulative distribution function of the folded Normal distribution. In particular,

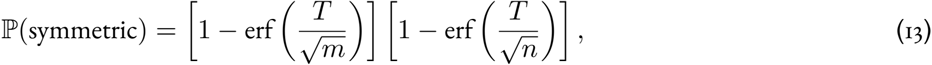

where erf is the error function. Since partitioning of A/B occurs independently of X/Y, for every given symmetric division S-S and D-D outcomes are equally likely. That is,

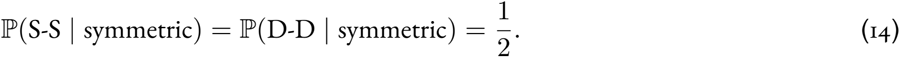

Thus,

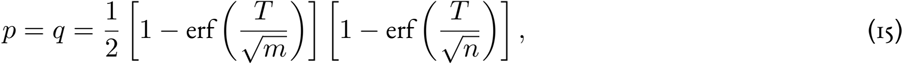

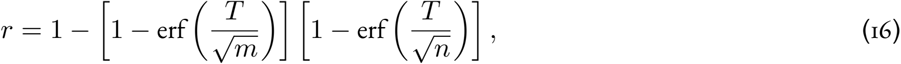

where *p, q* and *r* are defined in Eq. (1).

If 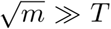 and 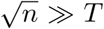 then these probabilities can be approximated by taking the leading order terms in the error function’s Maclaurin series, which gives

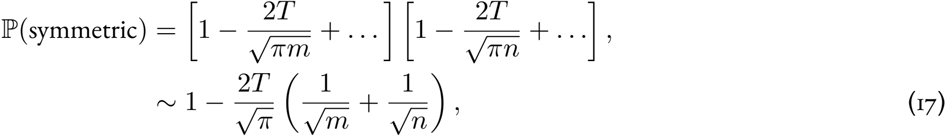

and so,

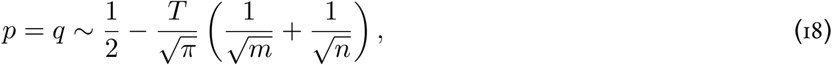

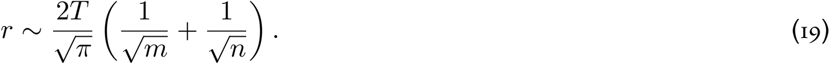

## References

[1] EA McCulloch, JE Till, Perspectives on the properties of stem cells. Nature Medicine 11, 1026–1028 (2005).

[2] IL Weissman, Stem cells: units of development, units of regeneration, and units in evolution. Cell 100, 157–168 (2000).

[3] G Martello, A Smith, The nature of embryonic stem cells. Annual Review of Cell and Developmental Biology 30, 647–675 (2014).

[4] BD Simons, H Clevers, Strategies for homeostatic stem cell self-renewal in adult tissues. Cell 145, 851–862 (2011).

[5] DT Scadden, The stem-cell niche as an entity of action. Nature 441, 1075–1079 (2006).

[6] G Mannino, et al., Adult stem cell niches for tissue homeostasis. Journal of Cellular Physiology 237, 239–257 (2022).

[7] CA Chacón-Martínez, J Koester, S. Wickström, Signaling in the stem cell niche: regulating cell fate, function and plasticity. Development 145, dev165399 (2018).

[8] KH Vining, DJ Mooney, Mechanical forces direct stem cell behaviour in development and regeneration. Nature Reviews Molecular Cell Biology 18, 728–742 (2017).

[9] N Pentinmikko, et al., Cellular shape reinforces niche to stem cell signaling in the small intestine. Science Advances 8, eabm1847 (2022).

[10] KB Blagoev, Organ aging and susceptibility to cancer may be related to the geometry of the stem cell niche. Proceedings of the National Academy of Sciences 108, 19216–19221 (2011).

[11] T Xin, V Greco, P Myung, Hardwiring stem cell communication through tissue structure. Cell 164, 1212–1225 (2016).

[12] BD MacArthur, A Ma’ayan, IR Lemischka, Systems biology of stem cell fate and cellular reprogramming. Nature Reviews Molecular Cell Biology 10, 672–681 (2009).

[13] MA Kinney, et al., A systems biology pipeline identifies regulatory networks for stem cell engineering. Nature Biotechnology 37, 810–818 (2019).

[14] E Fuchs, HM Blau, Tissue stem cells: architects of their niches. Cell Stem Cell 27, 532–556 (2020).

[15] X Lim, et al., Interfollicular epidermal stem cells self-renew via autocrine wnt signaling. Science 342, 1226–1230 (2013).

[16] C Parigini, P Greulich, Homeostatic regulation of renewing tissue cell populations via crowding control: stability, robustness and quasi-dedifferentiation. Journal of Mathematical Biology 88, 47 (2024).

[17] MD Johnston, CM Edwards, WF Bodmer, PK Maini, SJ Chapman, Mathematical modeling of cell population dynamics in the colonic crypt and in colorectal cancer. Proc. Natl. Acad. Sci. 104, 4008–4013 (2007).

[18] T Stiehl, A Marciniak-Czochra, Characterization of stem cells using mathematical models of multistage cell lineages. Mathematical and Computer Modelling 53, 1505–1517 (2011).

[19] G Bocharov, et al., Feedback regulation of proliferation vs. differentiation rates explains the dependence of CD4 T-cell expansion on precursor number. Proceedings of the National Academy of Sciences of the United States of America 108, 3318–3323 (2011).

[20] AD Lander, KK Gokoffski, FYM Wan, Q Nie, AL Calof, Cell lineages and the logic of proliferative control. PLoS Biology 7 (2009).

[21] T Alarcon, P Getto, A Marciniak-Czochra, M dM Vivanco, A model for stem cell population dynamics with regulated maturation delay. Conference Publications 2011, 32–43 (2011).

[22] T Stiehl, A Marciniak-Czochra, Stem cell self-renewal in regeneration and cancer: Insights from mathematical modeling. Current Opinion in Systems Biology 5, 112–120 (2017).

[23] X Chen, et al., Integration of external signaling pathways with the core transcriptional network in embryonic stem cells. Cell 133, 1106–1117 (2008).

[24] T Sato, et al., Single Lgr5 stem cells build crypt-villus structures in vitro without a mesenchymal niche. Nature 459, 262–265 (2009).

[25] L Conti, et al., Niche-independent symmetrical self-renewal of a mammalian tissue stem cell. PLoS Biology 3, e283 (2005).

[26] X Yin, et al., Niche-independent high-purity cultures of Lgr5+ intestinal stem cells and their progeny. Nature Methods 11, 106–112 (2014).

[27] AC Wilkinson, et al., Long-term ex vivo haematopoietic-stem-cell expansion allows nonconditioned transplantation. Nature 571, 117–121 (2019).

[28] CA Oedekoven, et al., Hematopoietic stem cells retain functional potential and molecular identity in hibernation cultures. Stem Cell Reports 16, 1614–1628 (2021).

[29] A Roshan, et al., Human keratinocytes have two interconvertible modes of proliferation. Nature Cell Biology 18, 145–156 (2016).

[30] H Shenghui, D Nakada, SJ Morrison, Mechanisms of stem cell self-renewal. Annual Review of Cell and Developmental Biology 25, 377–406 (2009).

[31] B Sunchu, C Cabernard, Principles and mechanisms of asymmetric cell division. Development 147, dev167650 (2020).

[32] JE Till, EA McCulloch, L Siminovitch, A stochastic model of stem cell proliferation, based on the growth of spleen colony-forming cells. Proceedings of the National Academy of Sciences 51, 29–36 (1964).

[33] T Suda, J Suda, M Ogawa, Disparate differentiation in mouse hemopoietic colonies derived from paired progenitors. Proceedings of the National Academy of Sciences 81, 2520–2524 (1984).

[34] T Suda, J Suda, M Ogawa, Single-cell origin of mouse hemopoietic colonies expressing multiple lineages in variable combinations. Proceedings of the National Academy of Sciences 80, 6689–6693 (1983).

[35] P Greulich, BD Simons, Dynamic heterogeneity as a strategy of stem cell self-renewal. Proceedings of the National Academy of Sciences 113, 7509–7514 (2016).

[36] P Greulich, BD MacArthur, C Parigini, RJ Sánchez-García, Universal principles of lineage architecture and stem cell identity in renewing tissues. Development 148, dev194399 (2021).

[37] A Wabik, PH Jones, Switching roles: the functional plasticity of adult tissue stem cells. The EMBO journal 34, 1164–1179 (2015).

[38] JA Knoblich, Mechanisms of asymmetric stem cell division. Cell 132, 583–597 (2008).

[39] BD MacArthur, IR Lemischka, Statistical mechanics of pluripotency. Cell 154, 484–489 (2013).

[40] AM Klein, BD Simons, Universal patterns of stem cell fate in cycling adult tissues. Development 138, 3103–3111 (2011).

[41] E Clayton, et al., A single type of progenitor cell maintains normal epidermis. Nature 446, 185–189 (2007).

[42] C Lopez-Garcia, AM Klein, BD Simons, DJ Winton, Intestinal stem cell replacement follows a pattern of neutral drift. Science 330, 822–825 (2010).

[43] DP Doupé, et al., A single progenitor population switches behavior to maintain and repair esophageal epithelium. Science 337, 1091–1093 (2012).

[44] AM Klein, T Nakagawa, R Ichikawa, S Yoshida, BD Simons, Mouse germ line stem cells undergo rapid and stochastic turnover. Cell Stem Cell 7, 214–224 (2010).

[45] HJ Snippert, et al., Intestinal crypt homeostasis results from neutral competition between symmetrically dividing Lgr5 stem cells. Cell 143, 134–144 (2010).

[46] C Parigini, P Greulich, Universality of clonal dynamics poses fundamental limits to identify stem cell self-renewal strategies. Elife 9, e56532 (2020).

[47] CS Potten, M Loeffler, Stem cells: attributes, cycles, spirals, pitfalls and uncertainties lessons for and from the crypt. Development 110, 1001–1020 (1990).

[48] NG Van Kampen, Stochastic processes in physics and chemistry. (Elsevier) Vol. 1, (1992).

[49] HW Fisher, J Yeh, Contact inhibition in colony formation. Science 155, 581–582 (1967).

[50] D Hanahan, RA Weinberg, The hallmarks of cancer. Cell 100, 57–70 (2000).

[51] Y Kitadate, et al., Competition for mitogens regulates spermatogenic stem cell homeostasis in an open niche. Cell Stem Cell 24, 79–92 (2019).

[52] RA Weinberg, The retinoblastoma protein and cell cycle control. Cell 81, 323–330 (1995).

[53] G Hateboer, RM Kerkhoven, A Shvarts, R Bernards, RL Beijersbergen, Degradation of E2F by the ubiquitinproteasome pathway: Regulation by retinoblastoma family proteins and adenovirus transforming proteins. Genes and Development 10, 2960–2970 (1996).

[54] F Hofmann, F Martelli, DM Livingston, Z Wang, The retinoblastoma gene product protects E2F-1 from degradation by the ubiquitin-proteasome pathway. Genes and Development 10, 2949–2959 (1996).

[55] LM Julian, A Blais, Transcriptional control of stem cell fate by E2Fs and pocket proteins. Frontiers in Genetics 6, 161 (2015).

[56] GD Amoutzias, DL Robertson, Y Van de Peer, SG Oliver, Choose your partners: dimerization in eukaryotic transcription factors. Trends in Biochemical Sciences 33, 220–229 (2008).

[57] NJ Marianayagam, M Sunde, JM Matthews, The power of two: protein dimerization in biology. Trends in Biochemical Sciences 29, 618–625 (2004).

[58] JD Klemm, SL Schreiber, GR Crabtree, Dimerization as a regulatory mechanism in signal transduction. Annual Review of Immunology 16, 569–592 (1998).

[59] Q Zhong, et al., Protein posttranslational modifications in health and diseases: Functions, regulatory mechanisms, and therapeutic implications. MedComm 4, e261 (2023).

[60] JM Lee, H. Hammarén, MM Savitski, SH Baek, Control of protein stability by post-translational modifications. Nature Communications 14, 201 (2023).

[61] SA Cuijpers, AC Vertegaal, Guiding mitotic progression by crosstalk between post-translational modifications. Trends in Biochemical Sciences 43, 251–268 (2018).

[62] H Cam, BD Dynlacht, Emerging roles for E2F: beyond the G1/S transition and DNA replication. Cancer cell 3, 311–316 (2003).

[63] JL Chong, et al., E2F1–3 switch from activators in progenitor cells to repressors in differentiating cells. Nature 462, 930–934 (2009).

[64] SM Rubin, J Sage, JM Skotheim, Integrating old and new paradigms of G1/S control. Molecular Cell 80, 183–192 (2020).

[65] SJ Morrison, J Kimble, Asymmetric and symmetric stem-cell divisions in development and cancer. Nature 441, 1068–1074 (2006).

[66] SP Acebron, E Karaulanov, BS Berger, YL Huang, C Niehrs, Mitotic Wnt signaling promotes protein stabilization and regulates cell size. Molecular Cell 54, 663–674 (2014).

[67] AB Alber, ER Paquet, M Biserni, F Naef, DM Suter, Single live cell monitoring of protein turnover reveals intercellular variability and cell-cycle dependence of degradation rates. Molecular Cell 71, 1079–1091 (2018).

[68] D Huh, J Paulsson, Random partitioning of molecules at cell division. Proceedings of the National Academy of Sciences 108, 15004–15009 (2011).

[69] J Parres-Gold, M Levine, B Emert, A Stuart, MB Elowitz, Principles of computation by competitive protein dimerization networks (2024).

[70] JM Paramio, C Segrelles, ML Casanova, JL Jorcano, Opposite functions for E2F1 and E2F4 in human epidermal keratinocyte differentiation. Journal of Biological Chemistry 275, 41219–41226 (2000).

[71] JL Vanderluit, et al., The Retinoblastoma family member p107 regulates the rate of progenitor commitment to a neuronal fate. Journal of Cell Biology 178, 129–139 (2007).

[72] A Balsalobre, J Drouin, Pioneer factors as master regulators of the epigenome and cell fate. Nature Reviews Molecular Cell Biology 23, 449–464 (2022).

[73] T Graf, T Enver, Forcing cells to change lineages. Nature 462, 587–594 (2009).

[74] H El-Samad, Biological feedback control—respect the loops. Cell Systems 12, 477–487 (2021).

[75] MA Muskavitch, Delta-Notch signaling and Drosophila cell fate choice. Developmental Biology 166, 415 (1994).

[76] D Sprinzak, et al., Cis-interactions between Notch and Delta generate mutually exclusive signalling states. Nature 465, 86 (2010).

[77] JA Goldman, KD Poss, Gene regulatory programmes of tissue regeneration. Nature Reviews Genetics 21, 511–525 (2020).

[78] I Sanidas, MS Lawrence, NJ Dyson, Patterns in the tapestry of chromatin-bound RB. Trends in Cell Biology 34, 288–298 (2024).

[79] A Blais, BD Dynlacht, E2f-associated chromatin modifiers and cell cycle control. Current opinion in cell biology 19, 658–662 (2007).

[80] P Greulich, (https://app.biorender.com/citation/676990f12651ac08361ca92a) (2024).

[81] R Rauber, C Cabreira, LB de Freitas, AC Turchetto-Zolet, M Margis-Pinheiro, The evolutionary history of the e2f and del genes in viridiplantae. Molecular Phylogenetics and Evolution 99, 225–234 (2016).

[82] T Berg, Modulation of protein–protein interactions with small organic molecules. Angewandte Chemie International Edition 42, 2462–2481 (2003).

[83] B Carter, K Zhao, The epigenetic basis of cellular heterogeneity. Nature Reviews Genetics 22, 235–250 (2021).

[84] P Cahan, GQ Daley, Origins and implications of pluripotent stem cell variability and heterogeneity. Nature Reviews Molecular Cell Biology 14, 357–368 (2013).

[85] K Hara, et al., Mouse spermatogenic stem cells continually interconvert between equipotent singly isolated and syncytial states. Cell Stem Cell 14, 658–672 (2014).

[86] L Ritsma, et al., Intestinal crypt homeostasis revealed at single-stem-cell level by in vivo live imaging. Nature 507, 362–365 (2014).

[87] SJ Ridden, HH Chang, KC Zygalakis, BD MacArthur, Entropy, ergodicity, and stem cell multipotency. Physical Review Letters 115, 208103 (2015).

[88] A Brunet, MA Goodell, TA Rando, Ageing and rejuvenation of tissue stem cells and their niches. Nature Reviews Molecular Cell Biology 24, 45–62 (2023).

[89] PS Stumpf, et al., Stem cell differentiation as a non-Markov stochastic process. Cell Systems 5, 268–282 (2017).

[90] J Paulsson, OG Berg, M Ehrenberg, Stochastic focusing: fluctuation-enhanced sensitivity of ntracellular regulation. Proceedings of the National Academy of Sciences 97, 7148–7153 (2000).

[91] W Bialek, Biophysics: searching for principles. (Princeton University Press), (2012).

[92] P Hänggi, Stochastic resonance in biology how noise can enhance detection of weak signals and help improve biological information processing. ChemPhysChem 3, 285–290 (2002).

[93] Y Arinobu, et al., Reciprocal activation of GATA-1 and PU.1 marks initial specification of hematopoietic stem cells into myeloerythroid and myelolymphoid lineages. Cell Stem Cell 1, 416–427 (2007).

[94] LA Boyer, et al., Core transcriptional regulatory circuitry in human embryonic stem cells. Cell 122, 947–956 (2005).

[95] OJ Rackham, et al., A predictive computational framework for direct reprogramming between human cell types. Nature Genetics 48, 331–335 (2016).

[96] X Liu, et al., Reprogramming roadmap reveals route to human induced trophoblast stem cells. Nature 586, 101–107 (2020).

[97] K Takahashi, S Yamanaka, Induction of pluripotent stem cells from mouse embryonic and adult fibroblast cultures by defined factors. Cell 126, 663–676 (2006).

[98] F. Müller, A Schuppert, Few inputs can reprogram biological networks. Nature 478, E4–E4 (2011).

[99] C Nerlov, E Querfurth, H Kulessa, T Graf, GATA-1 interacts with the myeloid PU.1 transcription factor and represses PU.1-dependent transcription. Blood 95, 2543–2551 (2000).

[100] A Reményi, et al., Crystal structure of a POU/HMG/DNA ternary complex suggests differential assembly of Oct4 and Sox2 on two enhancers. Genes & Development 17, 2048–2059 (2003).

[101] J Wang, et al., A protein interaction network for pluripotency of embryonic stem cells. Nature 444, 364–368 (2006).

[102] MA Skinnider, et al., An atlas of protein-protein interactions across mouse tissues. Cell 184, 4073–4089 (2021).

[103] L Velten, et al., Human haematopoietic stem cell lineage commitment is a continuous process. Nature Cell Biology 19, 271–281 (2017).

[104] C Weinreb, A Rodriguez-Fraticelli, FD Camargo, AM Klein, Lineage tracing on transcriptional landscapes links state to fate during differentiation. Science 367, eaaw3381 (2020).

[105] H Cheng, Z Zheng, T Cheng, New paradigms on hematopoietic stem cell differentiation. Protein & Cell 11, 34–44 (2020).

[106] HJ Beaumont, J Gallie, C Kost, GC Ferguson, PB Rainey, Experimental evolution of bet hedging. Nature 462, 90–93 (2009).

